# Early detection of ampicillin susceptibility in *Enterococcus faecium* with MALDI-TOF MS and machine learning

**DOI:** 10.1101/2025.07.14.664787

**Authors:** Thomas Pichl, Lucas Miranda, Nina Wantia, Karsten Borgwardt, Janko Sattler

**Affiliations:** Institute of Medical Microbiology, Immunology and Hygiene, TUM School of Medicine and Health, Department of Preclinical Medicine, Technical University of Munich, Germany; Department of Machine Learning and Systems Biology, Max Planck Institute of Biochemistry, Martinsried, Germany

## Abstract

**Background:** *Enterococcus faecium* can cause severe infections and is often resistant to the first-line antibiotic ampicillin. Consequently, clinicians usually prescribe broad-spectrum antibiotics, promoting the selection of multidrug-resistant bacteria. In this study, we investigate the application of machine learning techniques to detect ampicillin susceptibility directly from MALDI-TOF mass spectrometry. This technique could enable an earlier optimised treatment in infections with ampicillin-susceptible *E. faecium*.

**Methods:** Two datasets of clinical *E. faecium* MALDI-TOF spectra and their resistance phenotype were analysed: our own Technical University of Munich (TUM) dataset and the publicly available MS-UMG dataset. We tested logistic regression (LR) and LightGBM models on each dataset via nested cross-validation and explored transferability on the respective other dataset.

**Results:** LightGBM demonstrated slightly better performance than LR in identifying susceptible isolates in the TUM dataset (area under the precision-recall curve (AUPRC) 0.907 ± 0.016 vs 0.902 ± 0.030) as well as in the MS-UMG dataset (AUPRC 0.902 ± 0.029 vs 0.899 ± 0.054). External validation demonstrated good model transferability (AUPRC of 0.784 ± 0.039 when trained on MS-UMG; 0.804 ± 0.013 when trained on TUM). SHAP analysis consistently identified a top-ranked spectral feature corresponding to a peak at an m/z of 5091 in resistant isolate spectra.

**Conclusion:** This study demonstrates that LR and LightGBM models can identify ampicillin-susceptible *E. faecium* isolates from MALDI-TOF spectra and generalise well to unseen datasets. While clinical implementation currently still requires confirmatory testing, the addition of larger datasets in the future will support the development of more robust machine learning models.

## 1 Introduction

*Enterococcus faecium* is an important opportunistic pathogen that can cause severe infections such as endocarditis, hepatobiliary sepsis, and urinary tract infections. It is a Gram-positive bacterium within the *Enterococcaceae* family and part of the gastrointestinal microbiota [1]. While enterococcal species such as *E. faecalis* rarely exhibit resistance against ampicillin and vancomycin, *E. faecium* is a World Health Organisation-designated priority pathogen with multidrug-resistant strains being a global public health concern [2, 3].

Aminopenicillins such as ampicillin are generally well-tolerated and highly effective against susceptible *Enterococcus* strains. Mutations in the penicillin-binding protein 5 (PBP5) are considered the main cause of ampicillin resistance in *E. faecium*, though additional mechanisms remain less well characterised [4]. Due to the high prevalence of ampicillin resistance in healthcare-associated *E. faecium* isolates, glycopeptide antibiotics like vancomycin are commonly administered. However, the use of these broad-spectrum antibiotics promotes the selection and spread of resistant clones, e.g., *vanA-*mediated vancomycin-resistant *E. faecium*. Early identification of ampicillin-susceptible isolates would facilitate prompt targeted therapy and could therefore reduce unnecessary broad-spectrum antibiotic use and resistance dissemination.

Matrix-assisted laser desorption/ionization time-of-flight mass spectrometry (MALDI-TOF MS) is a commonly used method for bacterial identification [5]. In recent years, several studies investigated its potential for antimicrobial resistance prediction with machine learning (ML) models [6-8]. While most studies focused on *Staphylococcus aureus*, publications on *E. faecium* are scarce [9]. Public databases, such as the *MALDI-TOF Mass Spectra and Resistance Information on Antimicrobials from University Medical Center Göttingen* (MS-UMG), provide MALDI-TOF spectra and corresponding antimicrobial resistance data, facilitating research projects in this emerging field [10].

The objective of this study is to predict ampicillin susceptibility in *E. faecium* isolates using MALDI-TOF MS spectra in combination with ML. By analysing spectra derived from clinical samples at our microbiology laboratory at the Technical University of Munich (TUM), along with spectra provided by the MS-UMG dataset, we aim to develop ML models that can be seamlessly integrated into the routine workflows of diagnostic laboratories that already use MALDI-TOF MS for species identification.

## 2 Methods

### 2.1 Acquisition of spectra and susceptibility profiles

The TUM dataset comprises clinical MALDI-TOF spectra of *E. faecium* isolates collected at the TUM microbiology laboratory between August 2021 and May 2025 and the corresponding antimicrobial resistance profile. The majority of clinical samples originate from the TUM University Hospital Rechts der Isar and the TUM University Hospital German Heart Centre.

Spectral acquisition and bacterial identification were performed with the MALDI Biotyper® sirius one System (Bruker Daltonics, Bremen, Germany) using the manufacturer’s software solutions (flexcontrol 3.4, MBT Compass 4.1, MBT Compass Reference Library 2023). The direct transfer method was applied to bacterial isolates, followed by treatment with 70% formic acid (Merck, Darmstadt, Germany) and matrix application (α-Cyano-4-hydroxycinnamic acid, Bruker Daltonics, Bremen, Germany).

Antimicrobial resistance testing was conducted with the automated microdilution system VITEK® 2 (bioMérieux, Marcy-l’Étoile, France), disc diffusion tests (Becton Dickinson, Franklin Lakes, USA; MAST Group, Bottle, UK), or strip tests (Liofilchem, Roseto degli Abruzzi, Italy). *E. faecium* isolates were classified as ampicillin-resistant or -susceptible using the EUCAST breakpoints effective at the time of spectral acquisition. Only spectra with a MALDI-TOF score of 2.00 or higher and a single corresponding antibiogram were included.

*E. faecium* spectra and their corresponding antimicrobial susceptibility profiles were extracted from the MS-UMG dataset. We only included spectra from the diagnostic routine section of the database, matching the characteristics of the TUM dataset.

In both the TUM and MS-UMG datasets, spectra of isolates described as “susceptible, increased exposure (I)” in ampicillin susceptibility, as well as those with implausible antimicrobial susceptibility profiles, duplicate entries, and empty spectra were excluded from further analysis. Vancomycin-resistant *E. faecium* (VRE), which are generally ampicillin-resistant [11], were also excluded to avoid confounding.

### 2.2 Spectral preprocessing and feature alignment

Spectral preprocessing followed the protocol established by Weis et al. [6] employing the MALDIquant package version 1.22.2 [12] and included variance stabilization via square root transformation, signal smoothing using the Savitzky–Golay algorithm (half-window size: 10), baseline correction with 100 iterations of the SNIP algorithm, total ion current normalization, and trimming to the predefined m/z range of 2,000–20,000.

Spectral alignment was applied to correct for m/z shifts using a reference-based linear alignment approach adapted from publicly available code [13]. For internal validation, each dataset was aligned to its own reference peaks, defined as occurring in at least 90% of spectra within that dataset, with a tolerance of 0.2% m/z. For spectral alignment during external validation, the *external* dataset was aligned to the reference peaks of the *internal* training dataset. Aligned spectra were subsequently binned into 3 m/z windows, resulting in a feature vector of 6,000 dimensions per spectrum.

### 2.3 Model development and evaluation

The spectral feature vectors served as input for supervised ML models, with ampicillin susceptibility status as the dependent classification target (susceptible = positive class; resistant = negative class).

Two classification algorithms were employed in this study: LightGBM, a gradient boosting algorithm with strong performance in antibiotic resistance prediction from MALDI-TOF spectra [7], and logistic regression (LR), included for its interpretability and established use [9].

Models were developed using the scikit-learn (version 1.5.0) and lightgbm (version 4.4.0) Python libraries with hyperparameter tuning performed via RandomizedSearchCV. The Matthews Correlation Coefficient (MCC) was selected as the optimisation metric due to its suitability for imbalanced datasets.

Internal model performance was assessed using a stratified five-fold nested cross-validation approach (80/20 train/test split in the outer loop; 75/25 split on the inner loop). Performance metrics were computed and averaged across the outer folds, with macro means and standard deviations reported at a classification threshold of 0.5. Additionally, the area under the precision-recall curve (AUPRC) was calculated to provide a threshold-independent metric.

Model transferability was evaluated by training stratified five-fold cross-validation models on the MS-UMG and TUM datasets, with the respective other serving as the external test set. Results were averaged over five different random states to obtain reliable performance estimates.

SHAP (Shapley Additive exPlanations) values were calculated to determine the most relevant spectral features contributing to model predictions [14]. For further implementation details, please refer to the code in the supplemental material.

## 3 Results

### 3.1 Dataset characteristics

Extraction of MALDI-TOF MS spectra and corresponding susceptibility profiles of *E. faecium* isolates from the TUM dataset yielded 2109 spectra of ampicillin-resistant and 303 of susceptible isolates. The MS-UMG dataset comprises 1136 spectra of resistant isolates and 158 spectra of susceptible ones. Class distribution was similar in both datasets, with 14% and 12% susceptible isolates in the TUM and MS-UMG datasets, respectively.

### 3.2 5-fold nested cross-validation model performance on the TUM dataset

We first trained and evaluated a LR model using a 5-fold nested cross-validation approach on the TUM dataset. At a classification threshold of 0.5, the LR model showed good performance in determining ampicillin-susceptible isolates, achieving a positive predictive value (PPV) of 0.829 ± 0.048, a sensitivity of 0.878 ± 0.042, and an area under the precision-recall curve (AUPRC) of 0.902 ± 0.03 (**Table 1**). The model also performed strongly in identifying ampicillin-resistant isolates with a negative predictive value (NPV) of 0.982 ± 0.006 and a specificity of 0.973 ± 0.009.

**Table 1.** Average performance metrics (arithmetic mean) and standard deviations (SD) across folds for logistic Regression and LightGBM models using stratified 5-fold nested cross validation on the TUM dataset. All metrics, except for AUPRC (area under the Precision-Recall Curve), were calculated at a threshold of 0.5. PPV = Positive Predictive Value, NPV = Negative Predictive Value, MCC = Matthews Correlation Coefficient.

| Metric | Logistic regression |  | LightGBM |  |
| --- | --- | --- | --- | --- |
|  | Macro Mean | SD | Macro Mean | SD |
| PPV | 0.829 | 0.048 | 0.887 | 0.027 |
| Sensitivity | 0.878 | 0.042 | 0.855 | 0.067 |
| NPV | 0.982 | 0.006 | 0.979 | 0.010 |
| Specificity | 0.973 | 0.009 | 0.984 | 0.004 |
| MCC | 0.830 | 0.038 | 0.852 | 0.048 |
| AUPRC | 0.902 | 0.030 | 0.907 | 0.016 |

We then investigated whether model performance could be improved by employing LightGBM, as previously described [7]. Performance metrics of the LightGBM model showed a better PPV compared to LR with 0.887 ± 0.027, a slightly lower sensitivity of 0.855 ± 0.067, and a slightly higher AUPRC of 0.907 ± 0.016 (**Table 1**). Metrics for the detection of ampicillin-resistant isolates were comparable to LR.

### 3.3 5-fold nested cross-validation model performance on the MS-UMG dataset

To evaluate performance metrics of logistic regression and LightGBM on another, independent dataset, we performed nested cross-validation on the publicly available MS-UMG dataset.

While the LR model achieved a PPV of 0.781 ± 0.044 and a sensitivity of 0.811 ± 0.113, the LightGBM model showed a higher PPV (0.911 ± 0.092) but a lower sensitivity (0.671 ± 0.150) at the same classification threshold of 0.5 (**Table 2**). However, the threshold-independent AUPRC was 0.899 ± 0.054 for LR and slightly higher with 0.902 ± 0.029 for LightGBM, indicating strong performance in identifying ampicillin-susceptible isolates.

**Table 2.** Average performance metrics (arithmetic mean) and standard deviations (SD) across folds for logistic regression and LightGBM models using 5-fold nested cross validation on the MS-UMG dataset. All metrics, except for AUPRC (area under the Precision-Recall Curve), were calculated at a threshold of 0.5. PPV = Positive Predictive Value, NPV = Negative Predictive Value, MCC = Matthews Correlation Coefficient.

|  | *Logistic regression* |  | *LightGBM* |  |
| --- | --- | --- | --- | --- |
| **Metric** | **Macro Mean** | **SD** | **Macro Mean** | **SD** |
| PPV | 0.781 | 0.044 | 0.911 | 0.092 |
| Sensitivity | 0.811 | 0.113 | 0.671 | 0.150 |
| NPV | 0.974 | 0.015 | 0.956 | 0.019 |
| Specificity | 0.968 | 0.008 | 0.989 | 0.012 |
| MCC | 0.766 | 0.077 | 0.750 | 0.073 |
| AUPRC | 0.899 | 0.054 | 0.902 | 0.029 |

### 3.4 External model validation with LightGBM

We assessed model transferability by training LightGBM 5-fold cross-validation models on the TUM and MS-UMG datasets, with the respective other dataset serving as the test set across five different random states.

Both models demonstrated good overall transferability with an AUPRC of 0.784 ± 0.039 for the model trained on the TUM dataset and tested on the MS-UMG dataset, and an AUPRC of 0.804 ± 0.013 for the model trained on the MS-UMG dataset and tested on the TUM dataset (**Table 3**). PPV was high for the TUM-trained model (0.895 ± 0.089) at a classification threshold of 0.5, but sensitivity was low (0.410 ± 0.084). The opposite trend was observed for the MS-UMG trained model at the same threshold, which showed a sensitivity of 0.930 ± 0.012 and a PPV of 0.397 ± 0.017.

**Table 3.** Average performance metrics (arithmetic mean) and standard deviations (SD) across five random states for LightGBM models, using stratified 5-fold nested cross validation (80/20 train/test split) on the TUM dataset (training) and MS-UMG dataset (testing), and vice versa. All metrics, except for AUPRC (area under the Precision-Recall Curve), were calculated at a threshold of 0.5. PPV = Positive Predictive Value, NPV = Negative Predictive Value, MCC = Matthews Correlation Coefficient.

|  | *training TUM; testing MS-UMG* |  | *training MS-UMG; testing TUM* |  |
| --- | --- | --- | --- | --- |
| **Metric** | **Macro Mean** | **SD** | **Macro Mean** | **SD** |
| PPV | 0.895 | 0.089 | 0.397 | 0.017 |
| Sensitivity | 0.410 | 0.084 | 0.930 | 0.012 |
| NPV | 0.924 | 0.009 | 0.988 | 0.002 |
| Specificity | 0.992 | 0.009 | 0.797 | 0.013 |
| MCC | 0.569 | 0.034 | 0.529 | 0.019 |
| AUPRC | 0.784 | 0.039 | 0.804 | 0.013 |

### 3.5 Model interpretability via SHAP value analysis

In both LR and LightGBM internal validation models, SHAP value analysis of the ten most impactful spectral features (i.e., bins) from the outer folds’ test sets of the TUM dataset consistently identified a top-ranked feature in the m/z range of 5093-5096 (**Figure 1**). Interestingly, the same feature - or directly adjacent features – was also top-ranked in all internal LightGBM and LR models applied to the MS-UMG dataset. This trend persisted in the external validation models, where the feature was top-ranked across random states when trained on the TUM dataset and tested on the MS-UMG dataset and in four out of five random states when trained on MS-UMG and tested on TUM. All SHAP plots for internal and external model validation are provided in the supplementary material.

**Figure 1.**
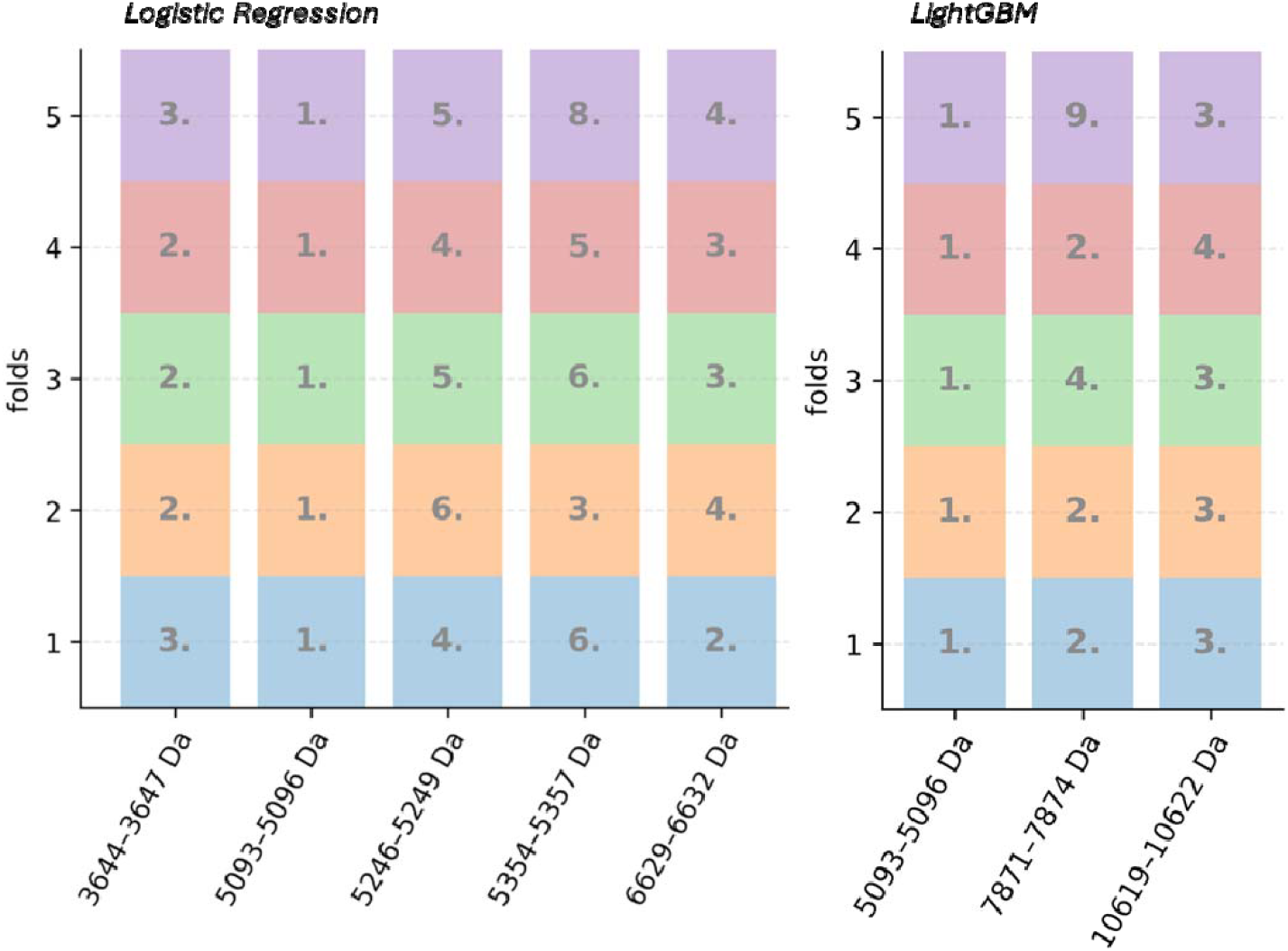
Top SHAP Features Predicting Ampicillin Susceptibility Across Folds. Mass-to-charge (m/z) ranges (x-axis) derived from the top ten most influential SHAP (SHapley Additive exPlanations) features of each fold in determining ampicillin susceptibility in *E. faecium* isolates from the TUM dataset, using stratified 5-fold nested cross-validation. Only features that appeared in the top ten SHAP ranking of all five folds are shown. Numbers within the boxes indicate the SHAP rank of the feature within the respective fold. Each fold is represented by a distinct colour and a corresponding number on the y-axis. Results are shown separately for logistic regression (left) and LightGBM (right).

Corresponding to the top-ranked feature at the m/z bin 5093-5096, we observed a distinct peak at an m/z of 5093.9 in the averaged spectra of unbinned, warped ampicillin-resistant *E. faecium* isolates from the TUM dataset. The peak was absent in the averaged spectra of susceptible isolates. By extrapolating from two conserved *E. faecium* ribosomal proteins in close proximity at m/z 4424 and 5353, we estimated the corresponding protein mass of the m/z 5093.9 peak at 5091 Da.

## 4 Discussion

Machine learning as a means to directly predict antimicrobial susceptibility from MALDI-TOF spectra has attracted growing interest in recent years [9]. In this context, *E. faecium* received comparatively little attention, and studies focused mostly on vancomycin resistance prediction [15, 16].

*E. faecium* is phylogenetically divided into clades which correlate with ampicillin susceptibility: Clade A1 (mostly resistant) is primarily found in healthcare settings, A2 (mostly susceptible) is linked to livestock, and B (also mostly susceptible) is frequently encountered in community environments [1, 17]. Faury et al. successfully differentiated *E. faecium* clades A1/A2 from clade B by constructing a MALDI-TOF spectra library but were unable to distinguish the 52 included isolates based on ampicillin susceptibility alone, with only 50% identified correctly [18]. In the present study, we set out to investigate ampicillin susceptibility prediction in *E. faecium* isolates from MALDI-TOF spectra derived from two clinical datasets - the TUM dataset and the MS-UMG dataset.

Clinical outcome data comparing ampicillin to glycopeptide antibiotics for ampicillin-susceptible enterococcal infections are scarce, with some retrospective studies reporting a lower mortality with ampicillin-based therapy [19, 20]. International guidelines recommend ampicillin, either as monotherapy or in combination, as the first-line treatment in severe infections caused by ampicillin-susceptible *E. faecium* [21]. Empiric treatment with glycopeptide antibiotics may be de-escalated once culture-based antibiotic susceptibility testing confirms ampicillin susceptibility.

Our model can support clinical decision-making in two potential scenarios. First, classification thresholds could be adjusted to maximise sensitivity, thereby increasing the chance of detecting ampicillin-susceptible isolates. This approach would allow for the early addition of ampicillin to empiric broad-spectrum antibiotic therapy, at the cost of accepting a higher number of false positives (lower PPV). This is analogous to the South Australia Health guidelines for empiric treatment of *Staphylococcus aureus* bloodstream infections, where cefazolin (optimal for oxacillin-susceptible *S. aureus)* and vancomycin (necessary for resistant strains) are both administered initially until susceptibility testing results become available [22]. Conversely, a classification threshold that prioritises the PPV may allow for earlier step-down from broad-spectrum antibiotics to ampicillin, reducing the unnecessary use of glycopeptide antibiotics. This approach accepts a lower sensitivity, whereby some susceptible isolates will be missed. However, with the current model performance as determined in this study, ampicillin susceptibility would still require confirmation via an additional, highly specific test. The precision-recall curves (supplementary **Figures S1 and S7**) highlight the trade-off between sensitivity and PPV in the context of predicting ampicillin-susceptibility.

SHAP value analysis revealed an adjusted m/z peak at 5091 as highly relevant for differentiating between the ampicillin-susceptible and -resistant classes, consistently across both datasets and different ML algorithms. Brackmann et al. identified oxidised bacteriocin T8 – also known as bacteriocin 43 or hiracin JM79 (Uniprot Q0Z8B6) – as the source of an m/z peak at 5092 in MALDI-TOF spectra of vancomycin-resistant *E. faecium* isolates using MALDI TOF/TOF (MS/MS) analysis [23]. Bacteriocin T8 presence has been shown to confer a competitive advantage relevant to the emergence of dominant *E. faecium* lineages [24]. Notably, a recent study found bacteriocin T8 to occur frequently in healthcare-associated *E. faecium* isolates, but rarely in community-acquired strains [25]. The spectral acquisition mode used by Brackmann et al. differed from the one used in this study, likely explaining the 1Da difference between the observed peaks. Whole-genome sequencing data were not available for the TUM or MS-UMG datasets; however, Faury et al. published genome sequences for 26 ampicillin-resistant and 26 -susceptible *E. faecium* isolates (Bioproject PRJEB56579) [18]. We detected bacteriocin T8 in 1 of 26 translated protein sequences from susceptible isolates and in 17 of 26 from resistant ones, based on alignments with ≥90% sequence identity and ≥90% query coverage. In summary, the peak observed in ampicillin-resistant *E. faecium* isolates in this study likely represents bacteriocin T8, which, while not directly mediating ampicillin resistance, may serve as a useful surrogate marker.

This study has several strengths. We used two independent clinical datasets from major German university hospitals, with spectra generated during routine diagnostic procedures. We assessed the performance of two machine learning algorithms -LightGBM and logistic regression-following established best practices to reduce overfitting and evaluate model interpretability. Finally, we demonstrated model transferability across datasets, evaluating broader diagnostic implementation.

However, there are important limitations. The retrospective design of this study limits its immediate applicability to prospective diagnostic workflows. The datasets used lacked genomic information and clade classification, preventing association between genes and the spectral peaks as well as clade-stratified analysis. Both datasets originate from German hospitals, and it remains uncertain whether similar results would be observed in other geographic settings with a different strain epidemiology.

Overall, our findings indicate good performance in the detection of ampicillin-susceptible *E. faecium* isolates using MALDI-TOF MS in combination with ML. This study contributes to the rapidly evolving field of machine learning for antibiotic susceptibility testing. In the future, the availability of additional and better characterised datasets will allow the development of more robust and generalisable models.

## Data availability

The Python and R scripts developed in this study are available on GitHub at https://github.com/MicrobeTom/E_faecium_ampicillin. MALDI-TOF spectra of *Enterococcus faecium* isolates used in this study are available from the Zenodo repository https://doi.org/10.5281/zenodo.15769315 distributed under a Creative Commons Zero v1.0 Universal licence.

## Declaration of Generative AI and AI-assisted technologies

ChatGPT (OpenAI) was used to assist with code generation and proofreading of the manuscript. All AI-assisted contributions were reviewed and verified by the authors.

## Ethical Approval

This study was reviewed by the TUM Ethics Committee (reference number 2024-543-W-SB), which determined that formal ethical approval was not required in accordance with Paragraph 15 of the Professional Code for Physicians in Bavaria (“Berufsordnung für Ärzte in Bayern”).

## Funding

None

## Competing interests

None declared

## Notes

https://zenodo.org/records/15769315

https://github.com/MicrobeTom/ECFMAmpi

### Competing Interest Statement

The authors have declared no competing interest.

### Summary of Updates

The discussion section has been updated to include a revised interpretation of the consistently top-ranked feature from the SHAP analysis.

