## Supplementary Materials for "Early detection of ampicillin susceptibility in *Enterococcus faecium* with MALDI-TOF MS and machine learning"

**1. Internal validation 5-fold nested cross-validation (TUM dataset)**

1.1. Logistic regression

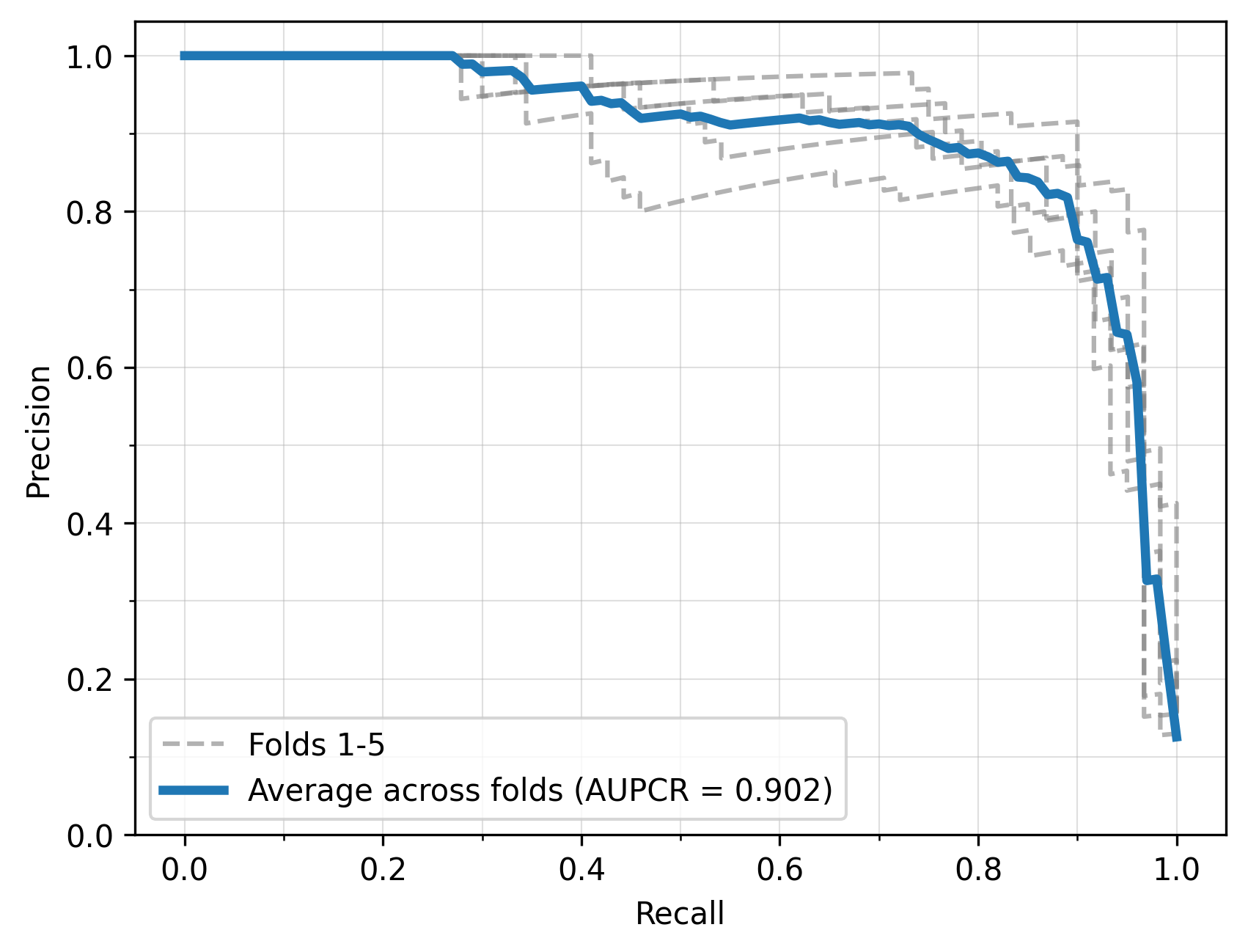

**Figure S1.** Precision–recall curves for all folds (gray dashed lines) and the average curve (arithmetic mean across folds, blue line) for the logistic regression model trained on the TUM dataset, using five-fold nested cross-validation. The area under the averaged precision–recall curve (AUPRC) is 0.902.

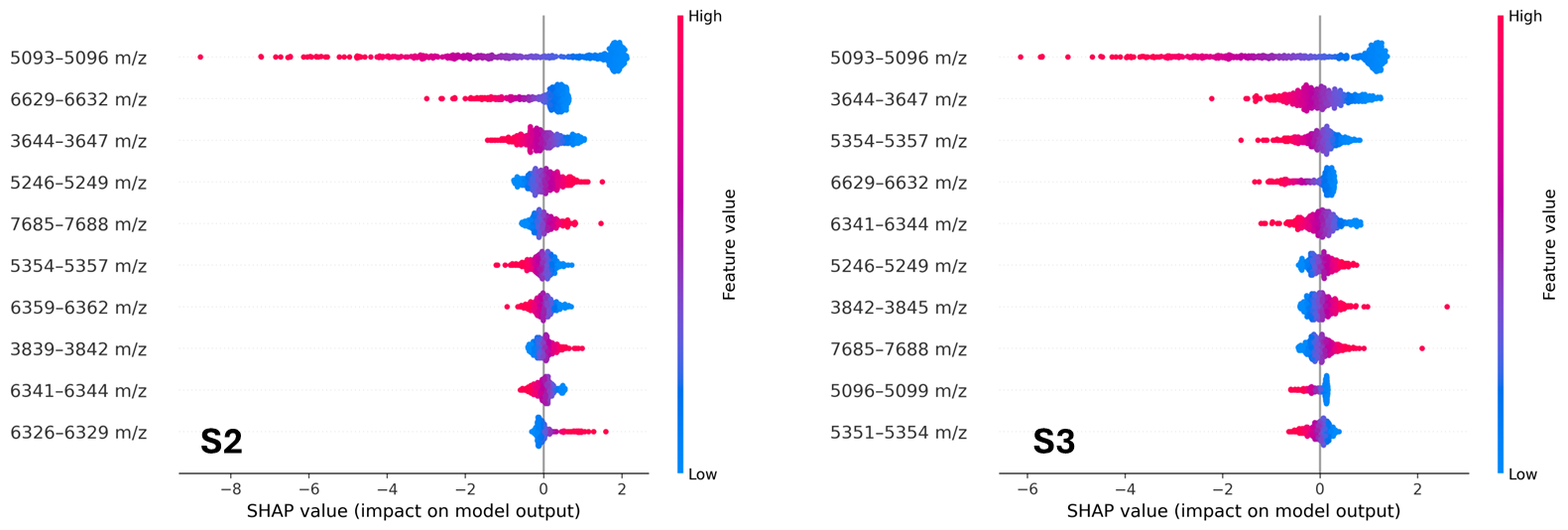

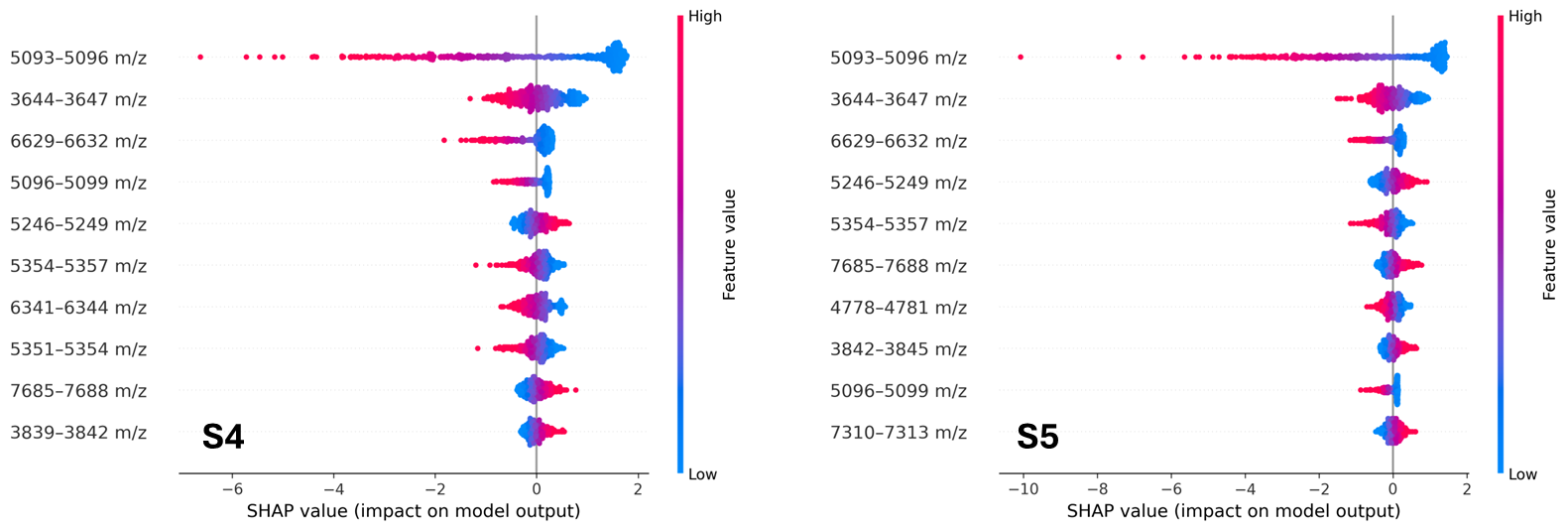

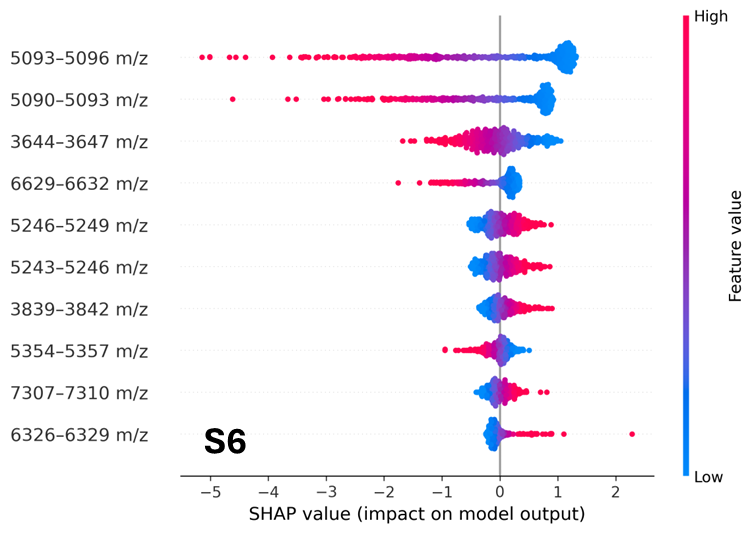

**Figures S2-S6.** SHAP (SHapley Additive exPlanations) beeswarm plots showing the ten most influential *E. faecium* spectral features for determining ampicillin susceptibility (susceptible = positive class; resistant = negative class) in the test sets of all five folds (in ascending order) from the stratified 5-fold nested cross-validation logistic regression model applied to the TUM dataset. Each dot corresponds to a single spectrum-feature pair, with the horizontal position indicating the SHAP value and its colour denoting the corresponding feature intensity (blue: low; red: high).

1.2 LightGBM

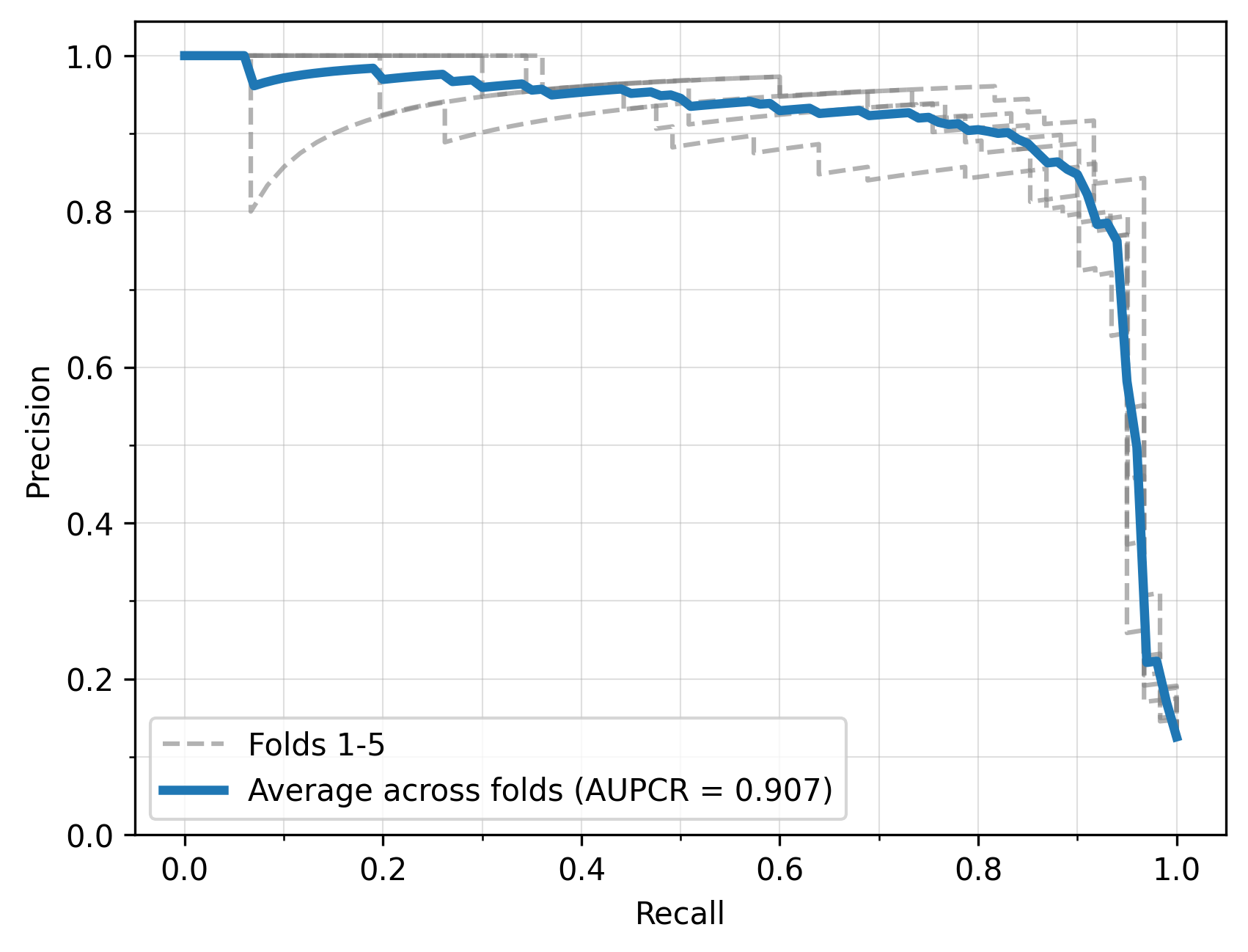

**Figure S7.** Precision–recall curves for all folds (gray dashed lines) and the average curve (arithmetic mean across folds, blue line) for the LightGBM model trained on the TUM dataset, using five-fold nested cross-validation. The area under the averaged precision–recall curve (AUPRC) is 0.907.

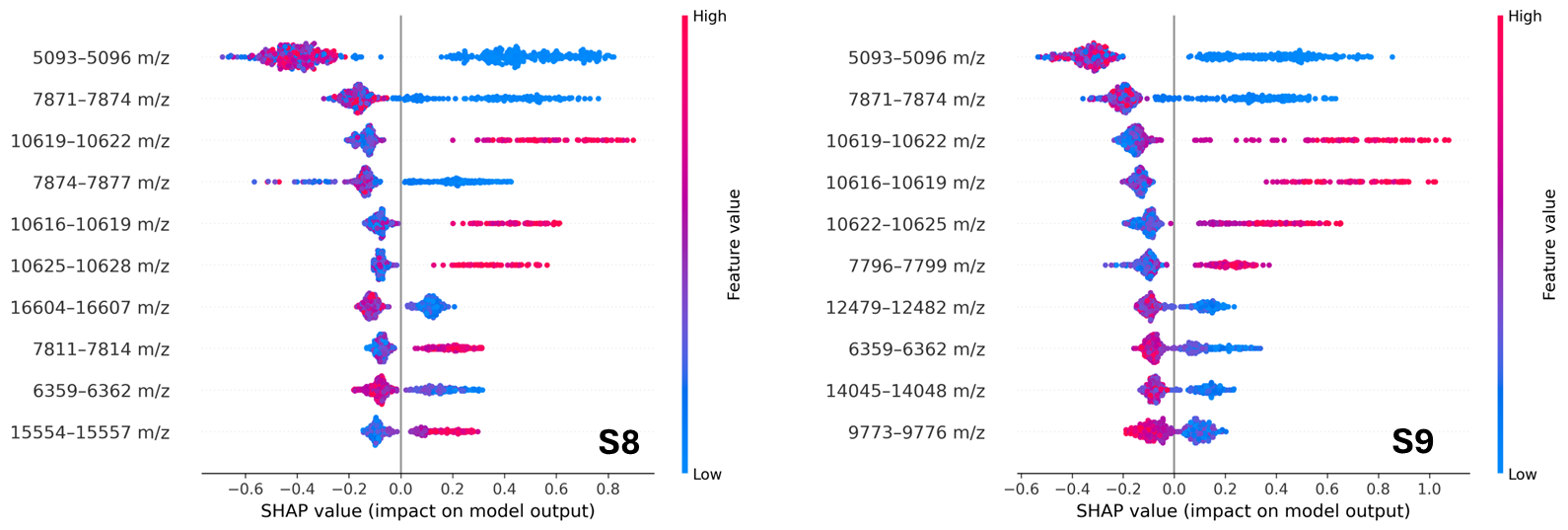

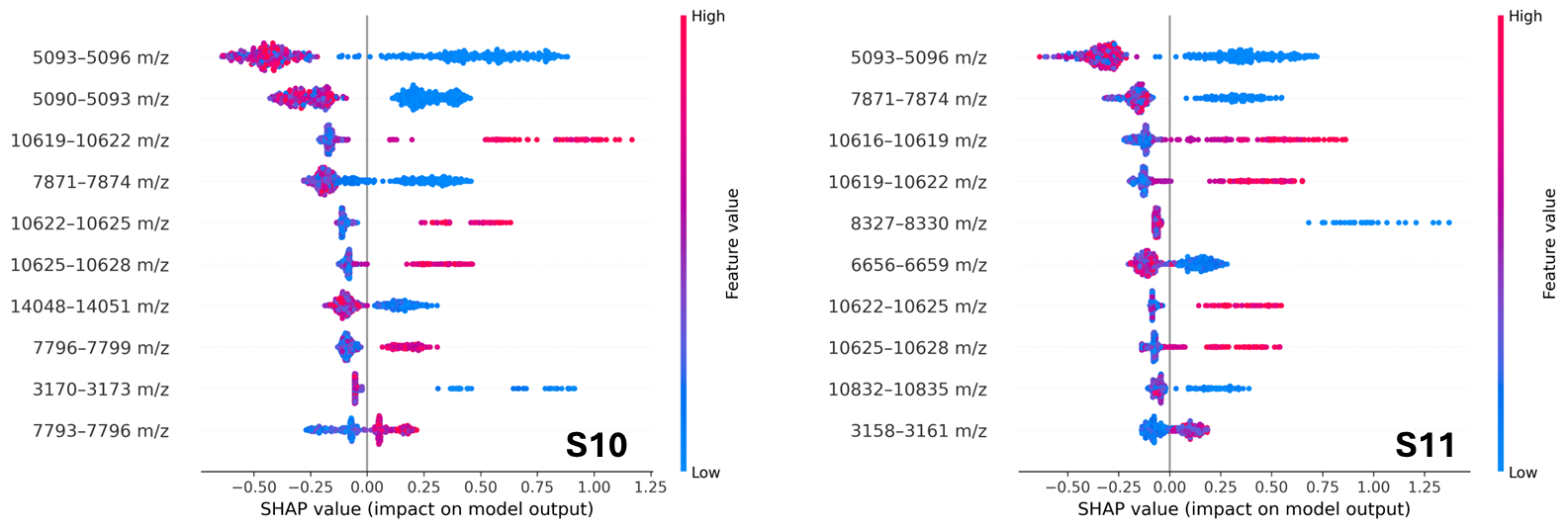

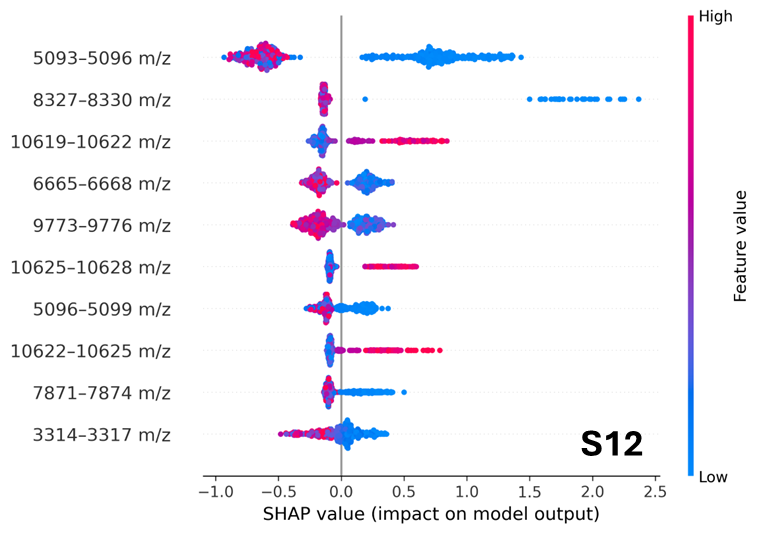

**Figures S8-S12.** SHAP (SHapley Additive exPlanations) beeswarm plots showing the ten most influential *E. faecium* spectral features for determining ampicillin susceptibility (susceptible = positive class; resistant = negative class) in the test sets of all five folds (in ascending order) from the stratified 5-fold nested cross-validation LightGBM model applied to the TUM dataset. Each dot corresponds to a single spectrum-feature pair, with the horizontal position indicating the SHAP value and its colour denoting the corresponding feature intensity (blue: low; red: high).

**2. Internal validation 5-fold nested cross-validation (MS-UMG dataset)**

2.1. Logistic regression

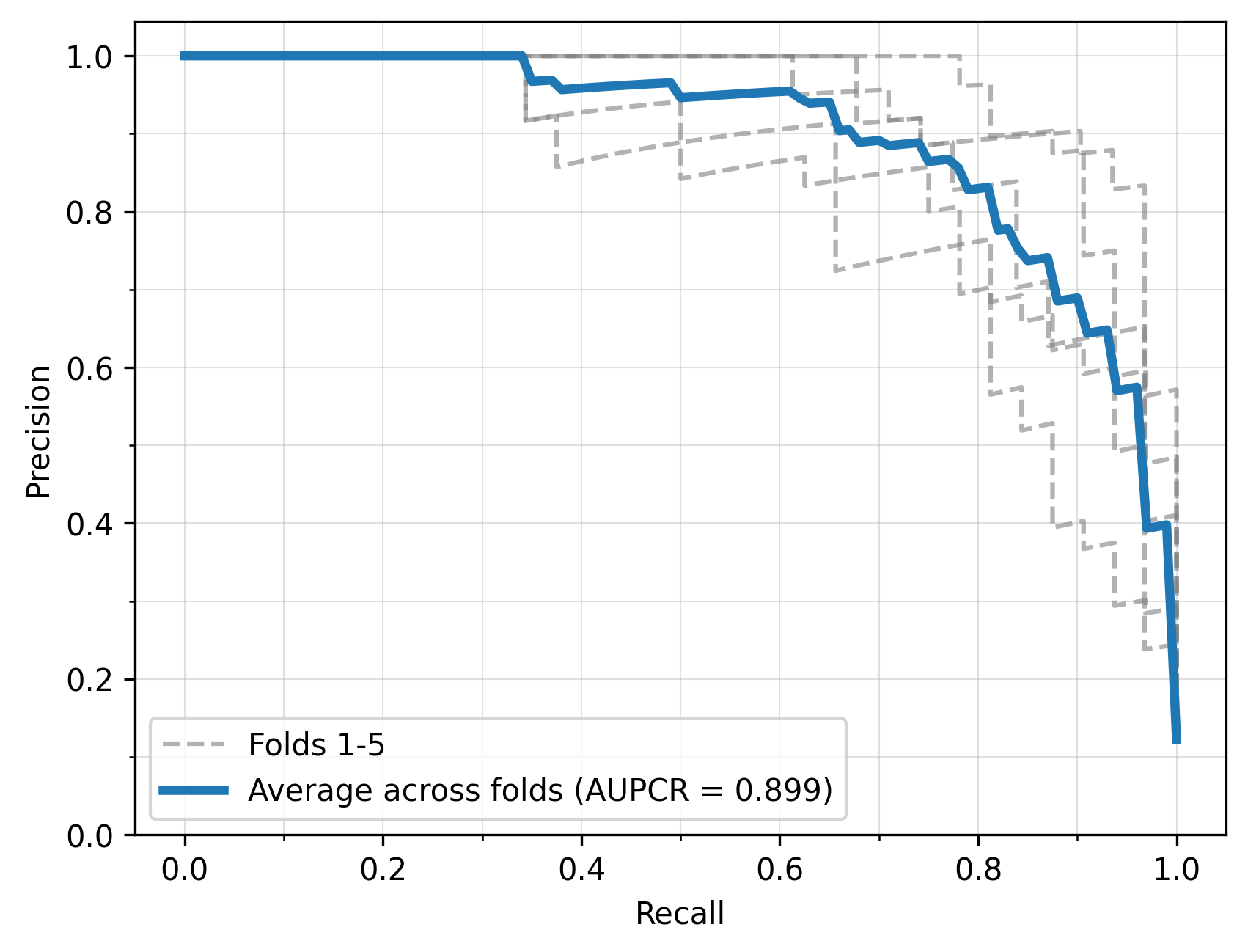

**Figure S13.** Precision–recall curves for all folds (gray dashed lines) and the average curve (arithmetic mean across folds, blue line) for the logistic regression model trained on the MS-UMG dataset, using five-fold nested cross-validation. The area under the averaged precision–recall curve (AUPRC) is 0.899.

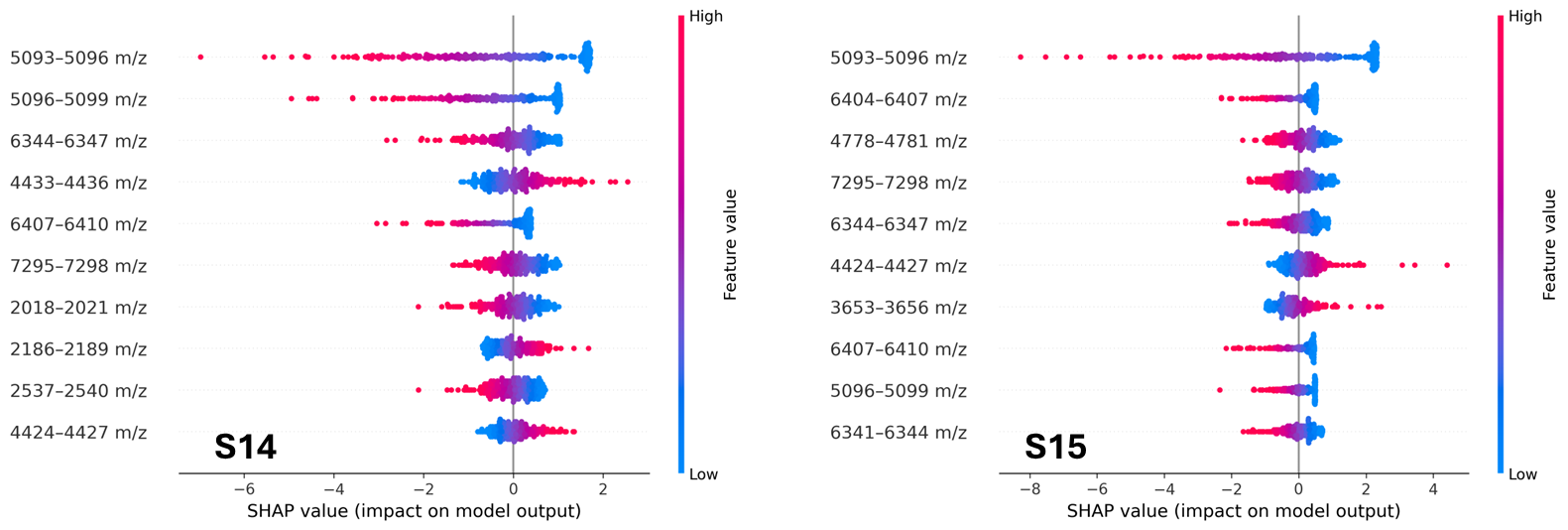

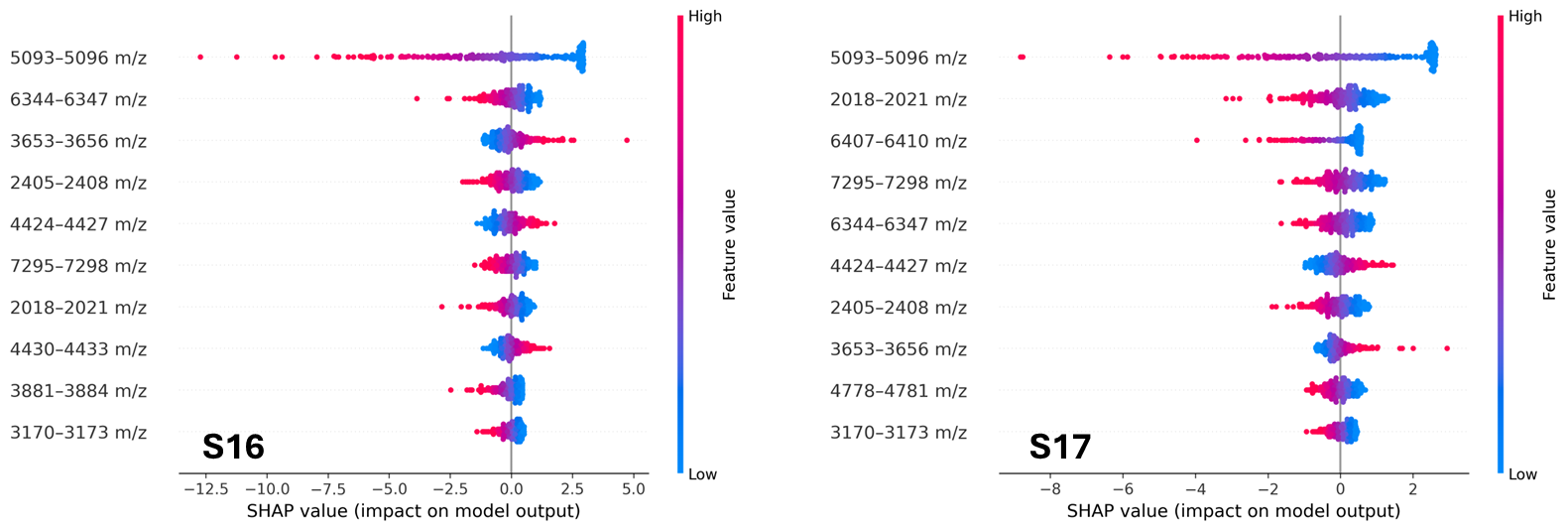

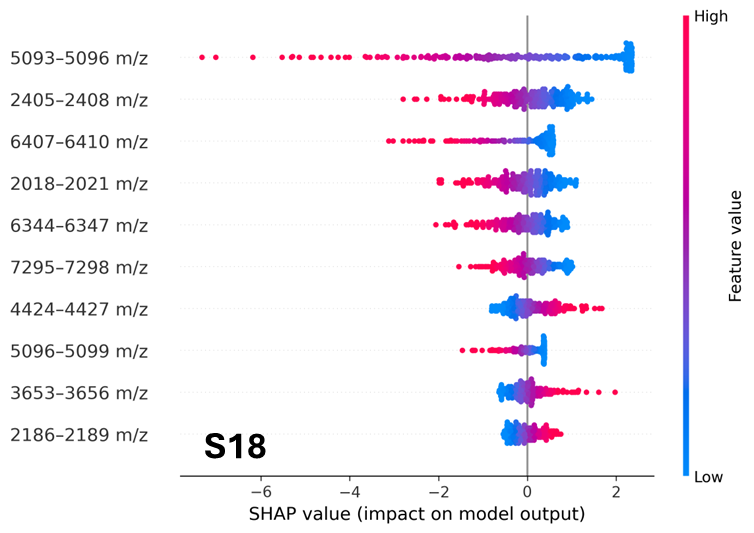

**Figures S14-S18.** SHAP (SHapley Additive exPlanations) beeswarm plots showing the ten most influential *E. faecium* spectral features for determining ampicillin susceptibility (susceptible = positive class; resistant = negative class) in the test sets of all five folds (in ascending order) from the stratified 5-fold nested cross-validation logistic regression model applied to the MS-UMG dataset. Each dot corresponds to a single spectrum-feature pair, with the horizontal position indicating the SHAP value and its colour denoting the corresponding feature intensity (blue: low; red: high).

2.2 LightGBM

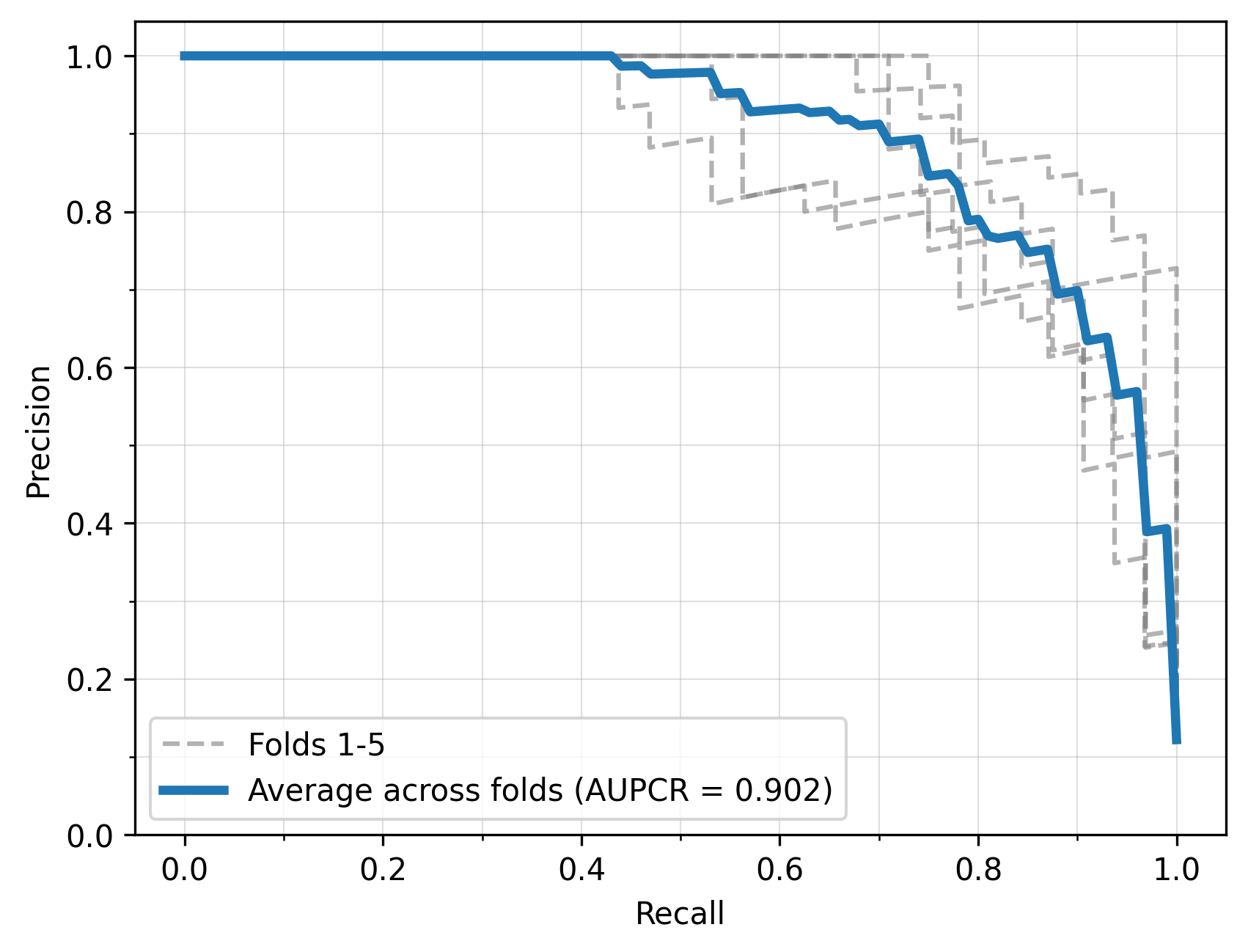

**Figure S19.** Precision–recall curves for all folds (gray dashed lines) and the average curve (arithmetic mean across folds, blue line) for the LightGBM model trained on the MS-UMG dataset, using five-fold nested cross-validation. The area under the averaged precision–recall curve (AUPRC) is 0.902.

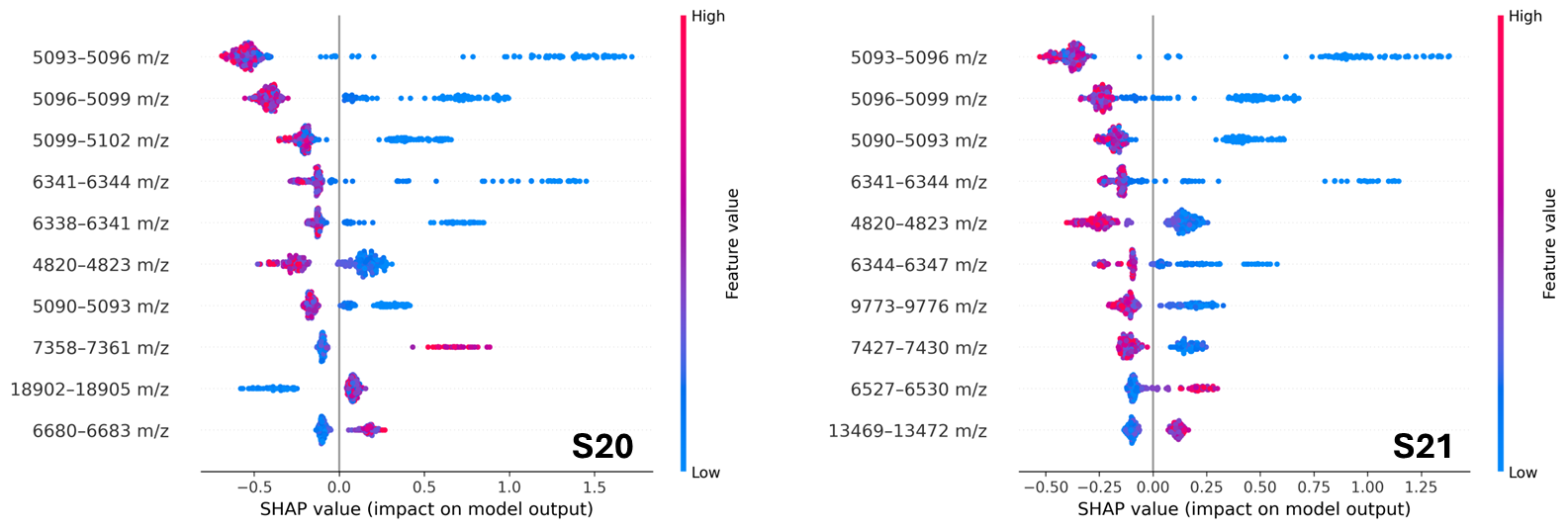

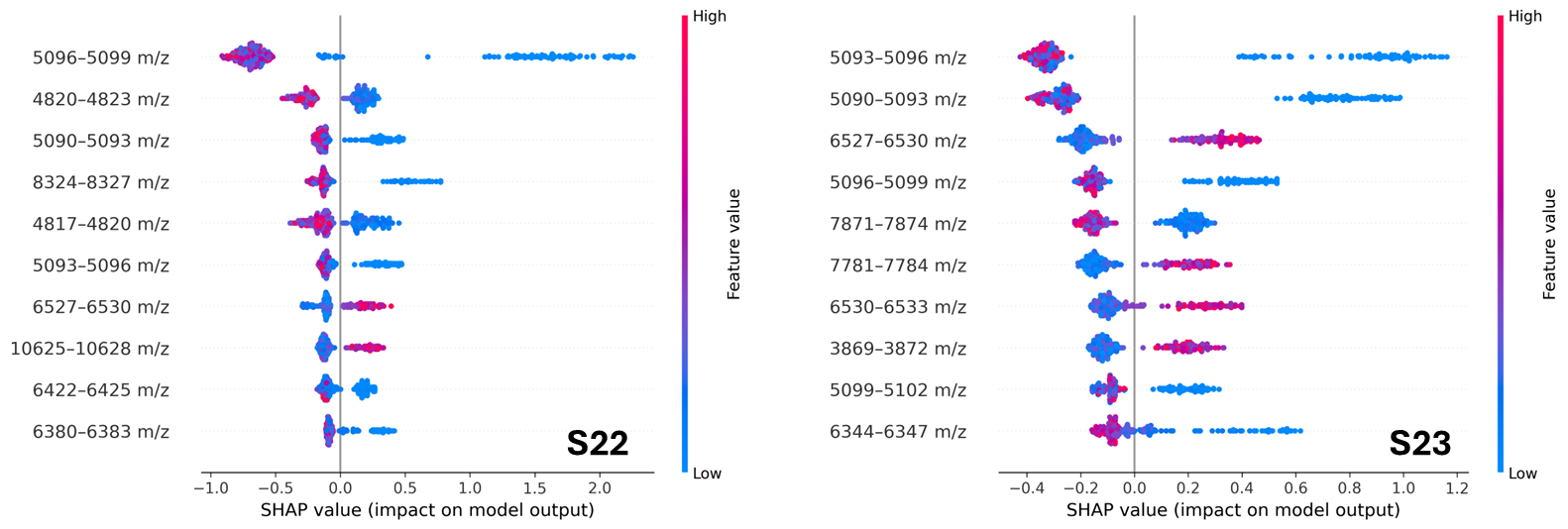

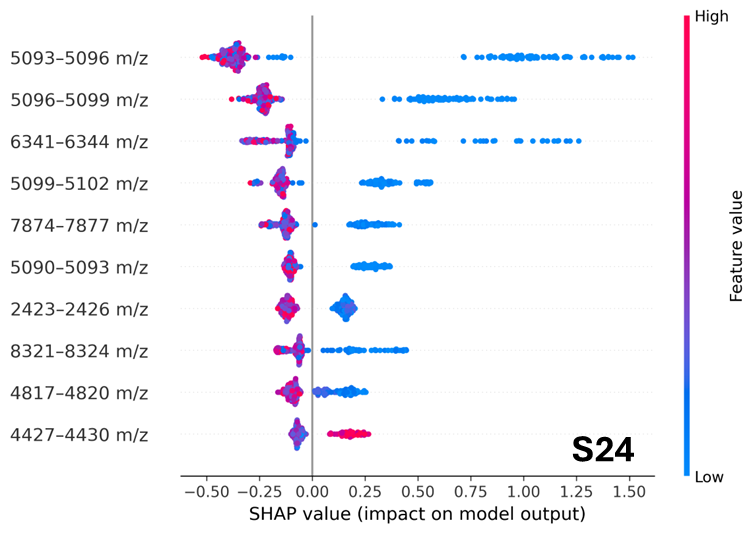

**Figures S20-S24.** SHAP (SHapley Additive exPlanations) beeswarm plots showing the ten most influential *E. faecium* spectral features for determining ampicillin susceptibility (susceptible = positive class; resistant = negative class) in the test sets of all five folds (in ascending order) from the stratified 5-fold nested cross-validation LightGBM model applied to the MS-UMG dataset. Each dot corresponds to a single spectrum-feature pair, with the horizontal position indicating the SHAP value and its colour denoting the corresponding feature intensity (blue: low; red: high).

**3. External 5-fold cross validation LightGBM model (trained on TUM; tested on MS-UMG)**

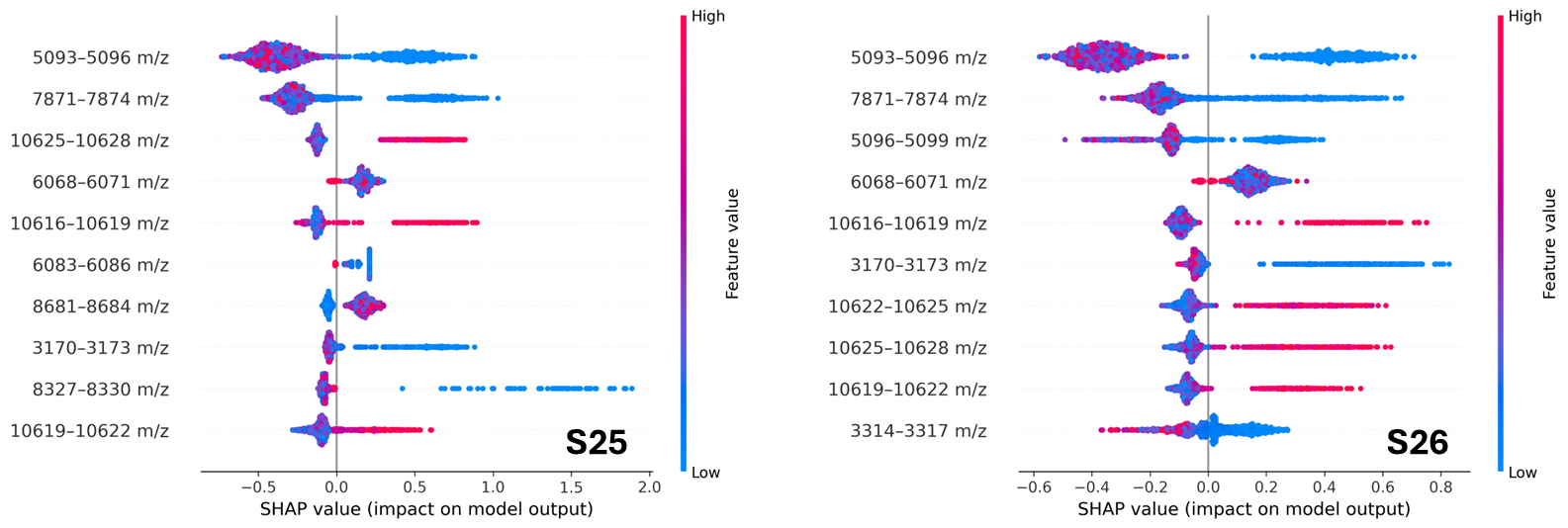

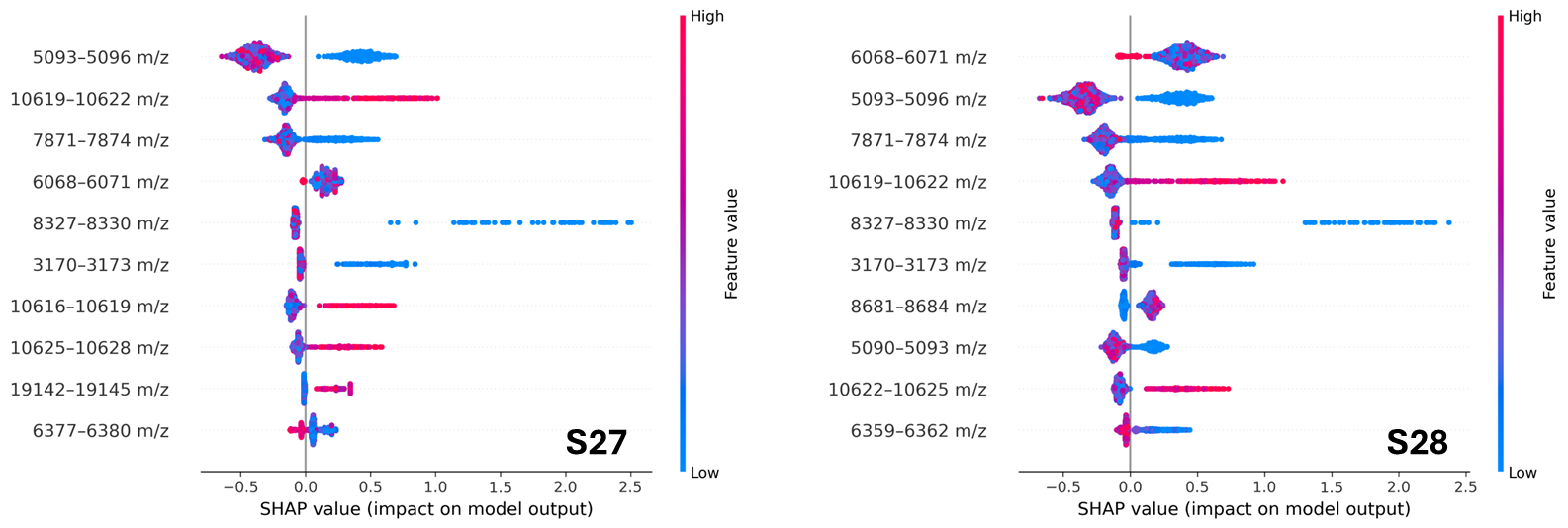

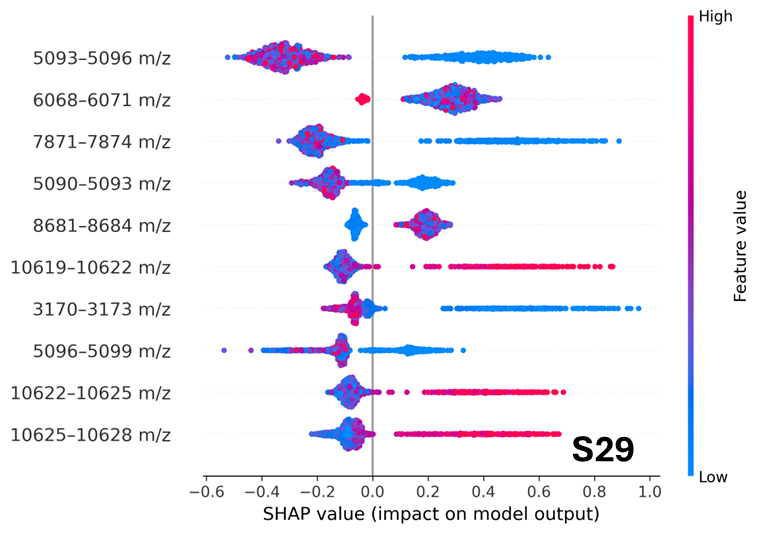

**Figures S25-S29.** SHAP (SHapley Additive exPlanations) beeswarm plots showing the ten most influential *E. faecium* spectral features for determining ampicillin susceptibility (susceptible = positive class; resistant = negative class) in the test sets of five random seeds from the 5-fold cross-validation LightGBM model trained on the TUM dataset and tested on the MS-UMG dataset. Each dot corresponds to a single spectrum-feature pair, with the horizontal position indicating the SHAP value and its colour denoting the corresponding feature intensity (blue: low; red: high).

**4. External 5-fold cross validation LightGBM model (trained on MS-UMG; tested on TUM)**

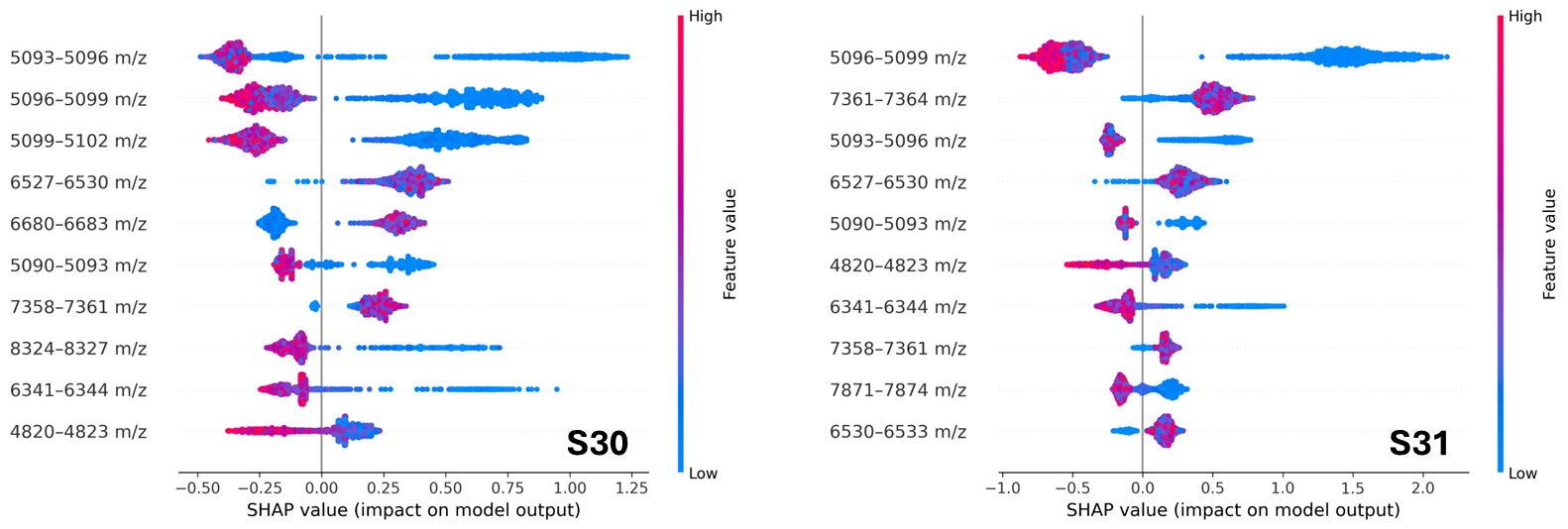

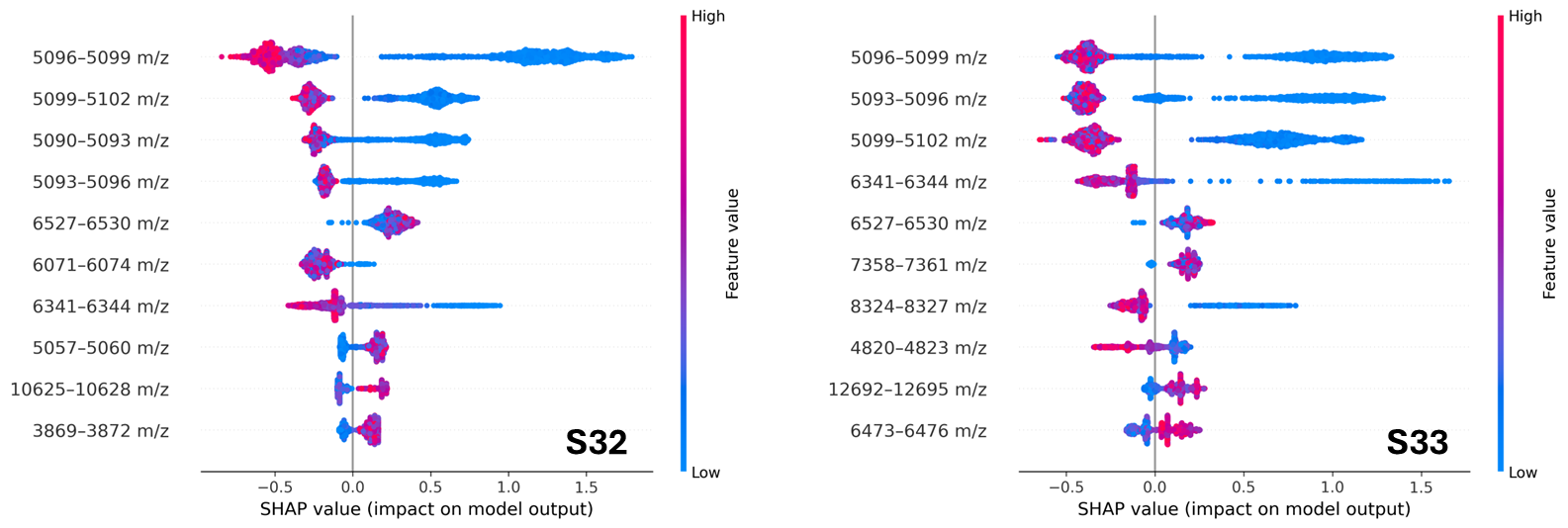

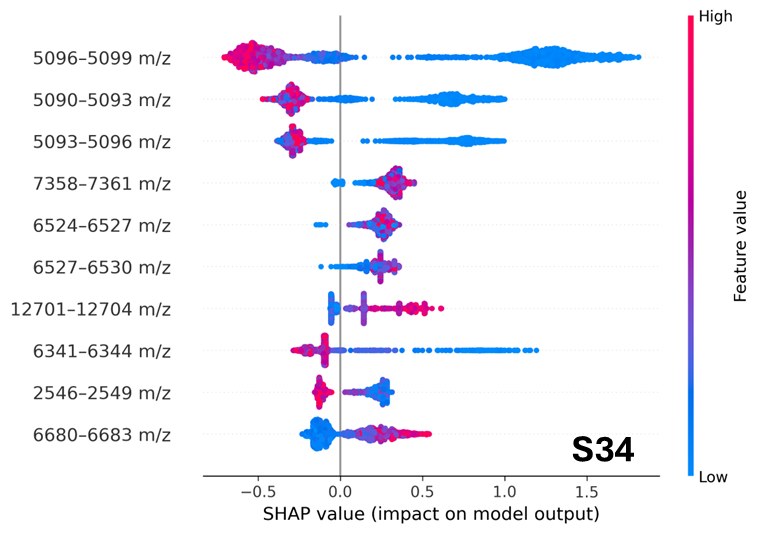

**Figures S30-S34.** SHAP (SHapley Additive exPlanations) beeswarm plots showing the ten most influential *E. faecium* spectral features for determining ampicillin susceptibility (susceptible = positive class; resistant = negative class) in the test sets of five random seeds from the 5-fold cross-validation LightGBM model trained on the MS-UMG dataset and tested on the TUM dataset. Each dot corresponds to a single spectrum-feature pair, with the horizontal position indicating the SHAP value and its colour denoting the corresponding feature intensity (blue: low; red: high).
